# Gene erosion and genome expansion in a group of highly host-specialized fungal phytopathogens

**DOI:** 10.1101/476267

**Authors:** Lamprinos Frantzeskakis, Márk Z. Németh, Mirna Barsoum, Stefan Kusch, Levente Kiss, Susumu Takamatsu, Ralph Panstruga

## Abstract

Due to their comparatively small genome size and short generation time, fungi are exquisite model systems to study eukaryotic genome evolution. Powdery mildew (PM) fungi present an exceptional case where their strict host dependency (a lifestyle termed obligate biotrophy) is associated with some of the largest fungal genomes sequenced so far (>100 Mbp). This size expansion is largely due to the pervasiveness of transposable elements (TEs), which can cover more than 70% of these genomes, and is associated with the loss of multiple conserved ascomycete genes (CAGs) required for a free-living lifestyle. To date, little is known about the mechanisms that drove this expansion, and information on ancestral PM genomes is lacking. We report the genome analysis of the early-diverged PM species *Parauncinula polyspora* that in contrast to most other PMs reproduces exclusively sexually. The *P. polyspora* genome is surprisingly small (<30 Mb) and sparsely equipped with TEs (<10%), despite the conserved absence of a common defense mechanism (RIP) involved in constraining repetitive elements. The genome still harbors the majority of the CAGs that are absent in the genomes of the recently evolved PMs. We speculate that TE spread might have been limited by its unique reproduction strategy and host features and further hypothesize that the loss of CAGs may promote the evolutionary isolation and host niche specialization of PM fungi. Limitations associated with this evolutionary trajectory might have been in part counteracted by the evolution of plastic, TE-rich genomes and/or the expansion of gene families encoding secreted virulence proteins.

## Introduction

Fungi provide a unique insight in the evolution of eukaryotic genomes due to their ubiquitous presence in diverse environments with different intensities of selection pressure (Gladieux et al. 2014). The genomes of phytopathogenic fungi in particular have been in the spotlight because of their peculiar genome architectures (Dong et al. 2015), which foster mechanisms that allow for the rapid adaptation to an ever-changing plethora of host resistance genes (Brown 2015). This high genome flexibility is considered to be a valuable tool for immune evasion, virulence and long-term survival (Raffaele and Kamoun 2012). Powdery mildews (PMs; Ascomycota, Erysiphales) are phytopathogens that exclusively colonize living host plants – a life-style termed obligate biotrophy (Spanu and Panstruga 2012). While some PMs have a broad host range and can infect members of different angiosperm families (Takamatsu 2013), others are restricted to a few host species belonging to the same plant genus or family (Troch et al. 2014). The best-known PM species are important pathogens of agriculturally and horticulturally relevant crops, while others are recognized as common pathogens of more than 10,000 angiosperms globally (Braun and Cook 2012). They typically propagate via short (several days long) asexual life cycles and the production of conidiospores, but can undergo sexual propagation by the formation of ascospores in particular circumstances (e.g. for overwintering; Glawe 2008). PMs also hold a special spot in filamentous plant pathogen genomics owing to the excessive size of their genomes, the vast amount of transposable elements (TEs) therein, and the large-scale loss of conserved fungal genes and associated cellular pathways (Bindschedler et al. 2016; Frantzeskakis et al. 2018; Spanu et al. 2010; Wicker et al. 2013). Recently, different laboratories have successfully managed to tackle technical challenges associated with the advanced genomic analysis of these pathogens, e.g. bottlenecks in extracting high molecular weight DNA (Feehan et al. 2017) or in assembling complex repetitive genomes to chromosome (arm) level (Frantzeskakis et al. 2018; Müller et al. 2018). Based on these methodological improvements, new information was provided on the population structure (Lu et al. 2016; Wicker et al. 2013), genome architecture (Frantzeskakis et al. 2018; Jones, L. et al. 2014; Müller et al. 2018; Wicker et al. 2013; Wu et al. 2018) and evolution (Menardo, Wicker et al. 2017) for some of the species and their specialized forms (formae speciales, ff.spp.). Based on the genomes of the barley and wheat PM pathogens, *Blumeria graminis* f.sp. *hordei* (*Bgh*) and f.sp. *tritici* (*Bgt*) respectively, it is suggested that TEs in powdery mildew genomes might be associated with the rapid turnover of virulence genes (effectors). This fast turnover can be recognized by the copy number variation of effector genes between the different specialized forms of *B. graminis* and their isolates (Frantzeskakis et al. 2018; Menardo, Praz et al. 2017; Müller et al. 2018). Additionally, it has been reported that PMs have a unique genomic architecture where TEs and genes are intertwined, while other typical attributes found in TE-inflated genomes are missing, e.g. AT-rich isochores or large-scale compartmentalization (Frantzeskakis et al. 2018; Müller et al. 2018). Since *B. graminis* has appeared relatively recently in the evolutionary history of PMs (Menardo, Wicker et al. 2017), the above-mentioned genome characteristics are also likely to represent a contemporary step in their evolution. In order to understand how the genomes of these pathogens evolved and what acted as a substrate for their unique genome architecture, it would be necessary to seek information in the genomes of early-diverged PM species. Here we present the genome of the PM species *Parauncinula polyspora*, pathogen of the East Asian oak tree *Quercus serrata*. It has been estimated that species of the genus *Parauncinula* have diverged from other genera of the *Erysiphales* 80-90 million years ago, rendering it one of the earliest diverged PM genera known to date (Takamatsu, Limkaisang et al. 2005; Takamatsu, Ninomi et al. 2005). The four species of *Parauncinula* (*P. curvispora, P. polyspora, P. septata* and *P. uncinata*) differ from other PMs in their unique morphology, host range, and also by lacking an asexual morph, i.e. conidiospores and conidiophores (Meeboon et al. 2017). Our analysis reveals that *P. polyspora* has a surprisingly compact genome with a substantially lower amount of TEs than the more recently evolved PMs, and this cannot be attributed to the presence of conserved ascomycete genome defense mechanisms such as repeat-induced point mutations (RIP). In addition, we report that the *P. polyspora* genome harbors a considerable amount of conserved ascomycete genes (CAGs) that were subsequently lost in other PM lineages. Taken together, the presented analysis of the *P. polyspora* genome gives unexpected insights into the evolutionary history of PM fungi and provides broad suggestions on how TE inflation can affect the genomes of fungal phytopathogens.

## Results

### The compact *P. polyspora* genome lacks large-scale compartmentalization and is indicative of homothallism

*Parauncinula polyspora* is believed to be an obligate biotroph that relies on living plant tissues for growth and reproduction (Takamatsu, Limkaisang et al. 2005), and in vitro cultures on artificial media of the species have never been reported. Therefore, we resorted in sampling infected *Q*. *serrata* leaves assumed to represent largely *P. polyspora* mycelium and ascomata (fruiting bodies), and sequenced the respective epiphytic metagenome. In order to avoid over-contamination, which could arise from sampling the entirety of the plant tissue, samples were prepared using cellulose acetate peelings (Both et al. 2005) of the leaf epiphytic microbial biome and the respective genomic DNA was subjected to short-read sequencing. As expected, the initial dataset contained sequences of a number of eukaryotic and prokaryotic taxonomic groups (Suppl. Figure 1A). Subsequently we assembled the respective short reads and followed a pipeline for stringent filtering to exclude both contaminating bacterial and eukaryotic sequences, assuming these are significantly less abundant than authentic *P. polyspora* sequences (Suppl. Figure 1B, Methods). After filtering for bacterial sequences, two major populations of scaffolds could be separated based on read depth. One of the two, with approximately 30x coverage for each of its 1321 scaffolds, contained sequences with similarity to PM fungi (average identity 60%, Suppl. Figure 1C), while the other (mostly with <5x coverage) contained a mix of additional fungal and plant sequences. Among the filtered contigs of the first population we identified a scaffold of extremely deep coverage (2384x) that is identical to the deposited nucleotide sequence of the internal transcribed spacer (ITS) region for the *P. polyspora* specimen voucher MUMH4928 (Suppl. Figure 2). DNA of this specimen was also sampled from PM-infected *Q. serrata* (Meeboon et al. 2017). We then annotated these scaffolds using a previously developed pipeline for the barley powdery mildew fungus (Frantzeskakis et al. 2018), and split them into 495 high and 826 low confidence contigs based on the relative frequency of Leotiomycete-related annotations along each sequence (Suppl. Figure 1B, Methods). The resulting 495 high confidence contigs contained 6046 genes in 28,01 Mb of sequence. Out of the annotated genes in these contigs, 97% have a detectable homolog in the Leotiomycetes (Suppl. Table 1). In terms of genome completeness, assayed using BUSCO (Simão et al. 2015), this assembly covers 90,75% of the common fungal gene space. Despite this percentage being somewhat lower compared to other sequenced PM genomes (Suppl. Table 2), it is surprisingly high for the biological material and procedure used, especially considering that earlier short-read-based approaches for other PM species (*Erysiphe pisi*, *Golovinomyces orontii*) have failed to provide assemblies with similarly high completeness scores (Spanu et al. 2010). Notably the ratio of single copy to duplicated BUSCO genes resembles that of other PM genome assemblies (Suppl. Table 2), indicating that our filtering method based on depth and sequence similarity has likely efficiently removed contaminating fungal sequences as well. For the downstream analysis, we used only the 495 high confidence contigs and the predicted annotations they contained. The assembled *P. polyspora* genome is very compact (28 Mb) compared to other sequenced PM species (Table 1), with a surprisingly low amount of repetitive elements (8,5%, Suppl. Table 3). In order to validate this observation we used a kmer-based approach (Marçais and Kingsford 2011) to calculate the genome size, which returned a similar estimate of 29,1 Mb. The difference in genome size to other sequenced PMs is also reflected by the length of the intergenic space, which is on average 3-4x smaller than e.g. in the case of *Bgh* (Figure 1A). Additionally, we noted that secreted protein-coding genes (SPs) do not constitute a separate compartment in the genome, i.e., there are no gene-rich/gene-sparse regions with an overrepresentation of SPs (Figure 1B, Suppl. Figure 3). Moreover, the comparatively few transposable elements (TEs), which nevertheless comprise representatives of all major groups (retrotransposons, LTR elements and DNA transposons; Suppl. Table 3), do not exhibit preferential insertion in the proximity of SPs (<1 kb, Figure 1C). Finally, we also found that in contrast to Bgh, the P. polyspora genome has a lower ratio of duplicated genes coding for SPs and non-SPs (Suppl. Table 4). We identified only a single scaffold harboring mating type genes (Suppl. Figure 4A), suggesting *P. polyspora* is a homothallic (self-fertile) PM species. These genes comprised a seemingly intact copy of MAT1-2-1 and a likely pseudogenized copy of MAT1-1-3 (Suppl. Figure 4B) residing ca. 10 kb away from each other on the same contig. However, we failed to identify a copy of MAT1-1-1, which is usually present in Leotiomycetes/PMs and in close physical proximity to MAT1-1-3 (Brewer et al. 2011; Frantzeskakis et al. 2018). Leotiomycete homologous sequences representing this gene were not only lacking in the 495 high confidence contigs but were also absent in the low confidence contigs and the contigs with lower coverage assumed to represent contaminations. Additionally, the gene SLA2, which is typically found near the MAT1 locus in many ascomycetes (Brewer et al. 2011), resides on a separate scaffold, flanked by genes that are non-syntenic to the canonical PM MAT1 locus. We also investigated the presence of single nucleotide polymorphisms (SNPs), and found that the majority of them are biallelic (98,2%, Suppl. Table 5). Considering the likely homothallic nature and taking into account that all PM genomes so far have been reported to be haploid during their vegetative growth phase, this finding indicates that our natural sample likely contained more than one *P. polyspora* isolate. To validate further the correct placement of this species as an early-diverged PM but also to corroborate the PM-related content of the high confidence contigs, we proceeded with generating a multi-type locus phylogeny based on 1964 single-copy orthologs of 16 sequenced Leotiomycetes (Figure 1D). The placement of the species by this approach at the base of the PM clade is in accordance with previous results based on ribosomal DNA (rDNA) sequences (Takamatsu, Limkaisang et al. 2005).

**Fig. 1.**
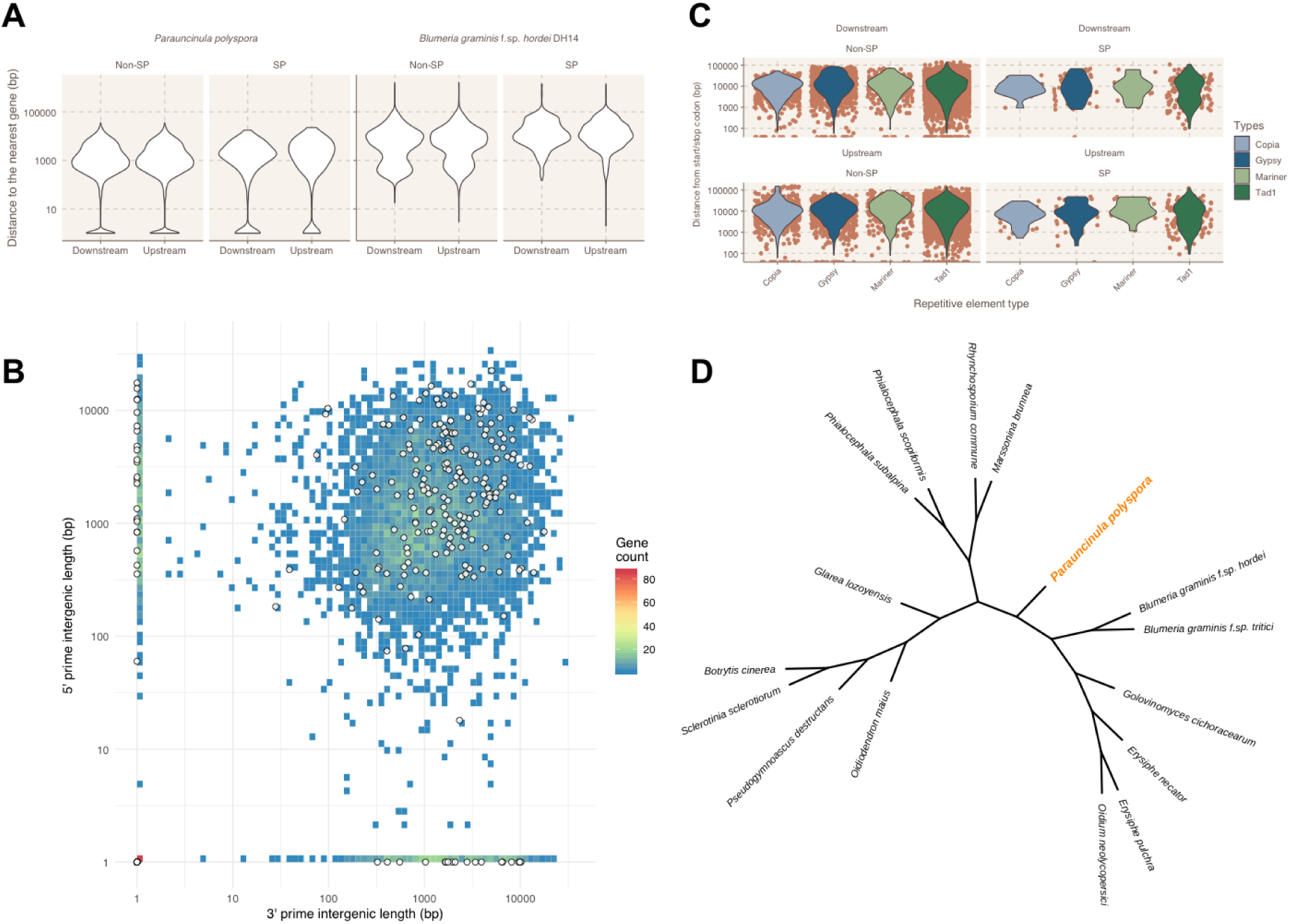
Characteristics of the *P. polyspora* genome. A) Violin plots illustrating the upstream (5′) and downstream (3′) intergenic length of the *P. polyspora* SP and non-SP-coding genes in comparison to *Bgh*. B) Gene density plot in relation to the 5’ and 3’ intergenic length for *P. polyspora*. White circles depict SP-coding genes. The number of genes for a given intergenic length is color-coded according to the indicated legend. C) Violin plots illustrating the distance of TEs from the transcriptional start/stop site of secreted and non-SP-coding genes. Orange dots indicate individual data points. D) Multi-locus phylogeny of selected Leotiomycete fungi based on 1964 single-copy orthologs identified by Orthofinder, was rendered by FastTree (ML Model: Jones-Taylor-Thornton) after alignment of the sequences with MAFFT.

**Table 1.**
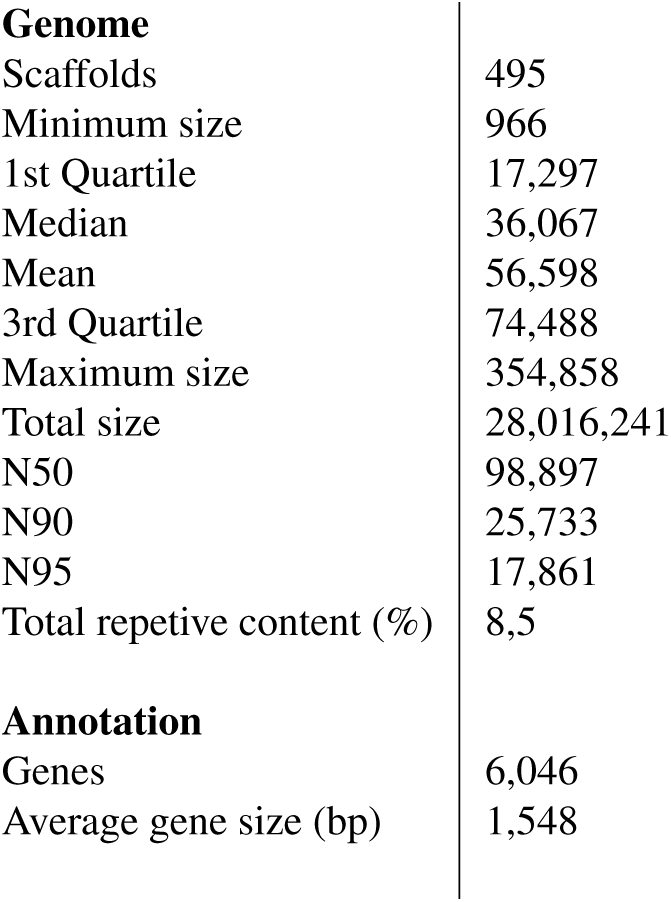
Genome assembly and annotation statistics

|  |  |
| --- | --- |
| **Genome** |  |
| Scaffolds | 495 |
| Minimum size | 966 |
| 1st Quartile | 17,297 |
| Median | 36,067 |
| Mean | 56,598 |
| 3rd Quartile | 74,488 |
| Maximum size | 354,858 |
| Total size | 28,016,241 |
| N50 | 98,897 |
| N90 | 25,733 |
| N95 | 17,861 |
| Total repetitive content (%) | 8,5 |
| **Annotation** |  |
| Genes | 6,046 |
| Average gene size (bp) | 1,548 |

### Genome expansion in the PM lineage cannot be solely attributed to the loss of the RIP machinery

We proceeded in examining whether the compactness of the *P. polyspora* genome is due to the presence of the genome defense mechanism of repeat-induced point mutation (Galagan and Selker 2004), which limits the spread of TEs and is absent in *Bgh* and other sequenced PM species (Jones, L. et al. 2014, 2014; Spanu and Panstruga 2012; Wu et al. 2018). We were unable to detect homologs of Masc1, Masc2, Rid-1 or Dim-2 (GenBank accessions AAC49849.1, AAC03766.1, XP_011392925.1 and XP_959891.1 respectively), which have been found to be associated with premeiotically-induced DNA methylation in *Ascobolus immersus* and *Neurospora crassa* (Freitag et al. 2002; Kouzminova and Selker 2001; Malagnac et al. 1997; Malagnac et al. 1999). Nevertheless, since these genes could have escaped the annotation process, or they could reside in genomic sequences which have either been removed during filtering or not been fully assembled, we additionally searched for genomic sequences that bear characteristic RIP signatures (i.e. overrepresentation of certain dinucleotide repeats). Initially we explored whether the *P. polyspora* genome has AT isochores, a typical feature of RIPed genomes such as in the case of *Leptosphaeria maculans* (Rouxel et al. 2011). We found that neither the *P. polyspora* nor the Bgh genome contains AT-rich isochores (Figure 2A). However, we observed that the intensity of the AT signature in genomes of other Leotiomycetes that contain the genes necessary for RIP varies (Suppl. Figure 5), with two exemplary cases for the presence of AT isochores being represented by *Marssonina brunnea* and *Rhynchosporium commune* (Figure 2A). We proceeded by calculating two indices for these four genomes that could be informative regarding the presence of RIP. These two indices (TpA/ApT and (CpA + TpG)/(ApC + GpT)) have been used previously in *N. crassa* to detect signatures of RIP in repetitive sequences (Margolin et al. 1998). They provide a measure for the prevalence and/or depletion of certain dinucleotides that are known results of RIP, while the respective baseline frequencies are calculated from non-repetitive genomic sequences of the same genome. In the case of *N. crassa*, sequences with an TpA/ApT index higher than 0,89 and/or an (CpA + TpG)/(ApC + GpT) index lower than 1,03 suggest a biased frequency of AT dinucleotides caused by RIP (Margolin et al. 1998). The baseline frequencies may vary between different species, as expected by the different overall nucleotide frequency of their genomes. In the genomes of *Bgh* and *P. polyspora*, which seemingly lack RIP-related genes (see above), these indices likewise point to an absence of RIP. By contrast, respective values for *M. brunnea* and *R. commune* indicate the presence of RIPed sequences, which is in agreement with the presence of AT-rich isochores and the genes of the RIP machinery in these genomes (Figure 2B, C). Repetitive sequences that deviate from the set thresholds in *Bgh* and *P. polyspora* could be relics of ancient RIP events, as it has been previously suggested (Amselem et al. 2015). Interestingly, the majority of the TEs in *P. polyspora* show high nucleotide sequence divergence (30-40%; Figure 2D), in contrast to *Bgh* where recent TE bursts have been observed, resulting in highly similar (>90% sequence identity) TE copies (Frantzeskakis et al. 2018).

**Fig. 2.**
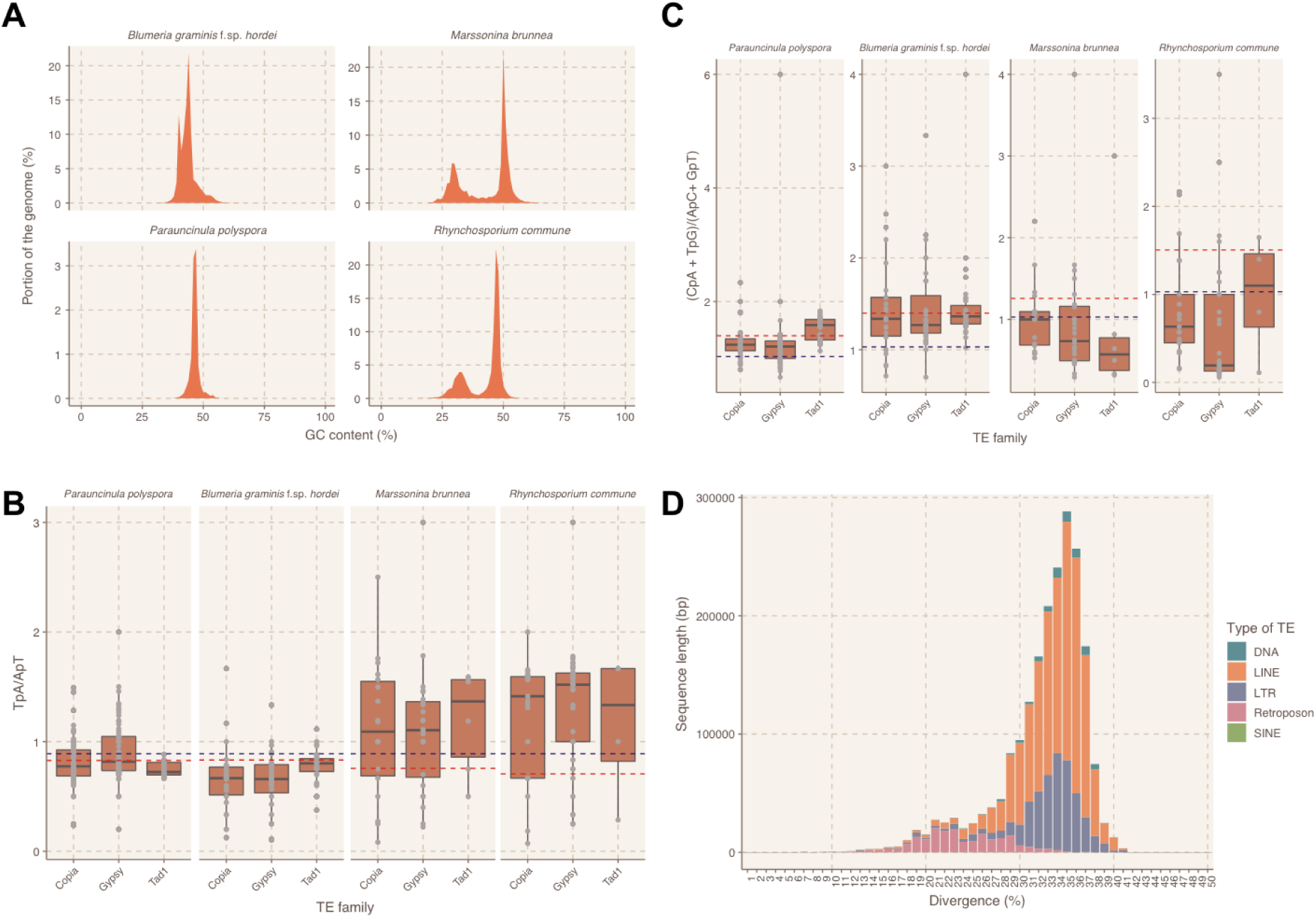
Analysis for signatures of RIP in the *P. polyspora* and related Leotiomycete genomes. A) GC content profile of the *P. polyspora, Bgh, M. brunnea* and *R. commune* genomes. B) and C) RIP index analysis for repetitive sequences of the four genomes. The blue line depicts the thresholds set from N. crassa (0.89 and 1.03 respectively; Margolin et al. 1998), while the red line indicates the threshold values obtained by the non-TE containing genomic sequences of each respective genome. In B, the TpA/ApT ratio is given, in C the (CpA + TpG)/(ApC + GpT) ratio is shown. D) Divergence analysis for the TEs in *P. polyspora* genome. The histogram illustrates the total sequence length (in bp) with a given nucleotide sequence divergence (in %) for different types of TEs according to the color code in the legend.

### Powdery mildew genomes exhibit a lineage-specific loss of conserved ascomycete genes

Next, we sought to determine if a set of conserved ascomycete genes (CAGs) that were previously found to be missing in the genome of the barley PM fungus (Spanu et al. 2010) could be identified in the annotation of the *P. polyspora* genome. In addition, we surveyed additional PM, Leotiomycete and ascomycete proteomes to reevaluate the conservation of these proteins throughout the ascomycete lineage. Interestingly a major portion of these genes (77 out of 98) could be detected in the *P. polyspora* genome (Figure 3A), in contrast to other PMs where the presence of these genes is low but somewhat variable. For example, in the *Bgh* genome only 17 out of 98 CAGs were found. The presence of 17 genes that were previously considered to be absent (Spanu et al. 2010) might be explained by the improved assembly and annotation of the *Bgh* reference genome (Frantzeskakis et al. 2018). Some of the genes found to be present in *P. polyspora* are critical for common biochemical pathways such as glutathione metabolism (Figure 3A). However, the preservation of genes in *P. polyspora* does not apply to all otherwise widely conserved functional modules, as for example the RIP mechanism is lacking (see above) and the number of carbohydrate-active enzymes (CAZymes), which are abundant in close relatives of PMs (e.g. in *Botrytis cinerea* and *Sclerotinia sclerotiorum*, is comparatively low (Figure 3B, Suppl. Table 6). Notably, the results of this analysis indicate that the loss of some of these conserved genes might have happened independently for the different PM sub-lineages (e.g. for the homologs of YHL016C and YIR023W or for YGL202W), while others were seemingly lost prior to the diversification of the PMs (e.g. for the homologs involved in glutamate metabolism). We also manually inspected the position of the respective *P. polyspora* orthologs in the associated phylogenetic trees generated by Orthofinder to reduce the likelihood of contaminations from other fungi that might have persisted through the filtering step, possibly indicating a false-positive presence of the respective genes (Suppl. Figure 6). In addition, we found that beyond a number of unique PFAM-annotated functional domains that can be detected in the genome *P. polyspora* when compared to *Bgh* (Suppl. Table 7), there is a considerable fold difference (>3) in the presence of 27 PFAM domains in *P. polyspora* when compared to *Bgh* (Figure 3C). Particularly noteworthy in this context are the flavin-binding monooxygenase-like and glucose-methanol-choline (GMC) oxidoreductase functional domains, which each show a >8-fold increased presence in relation to *Bgh* (Figure 3C). By contrast, in comparison to the *Bgh* genome the *P. polyspora* genome appears to be depleted for genes encoding members of the peptidase family S41 (>10-fold lower content than in Bgh; Figure 3C). This comparison also emphasizes some of the unique aspects of the *Bgh* genome, such as the expansion of genes encoding proteins with Sgk2 domains (SUPERFAMILY SSF56112; Kusch et al. 2014), which cannot be observed to the same extent in the other PM genomes (Suppl. Figure 7).

**Fig. 3.**
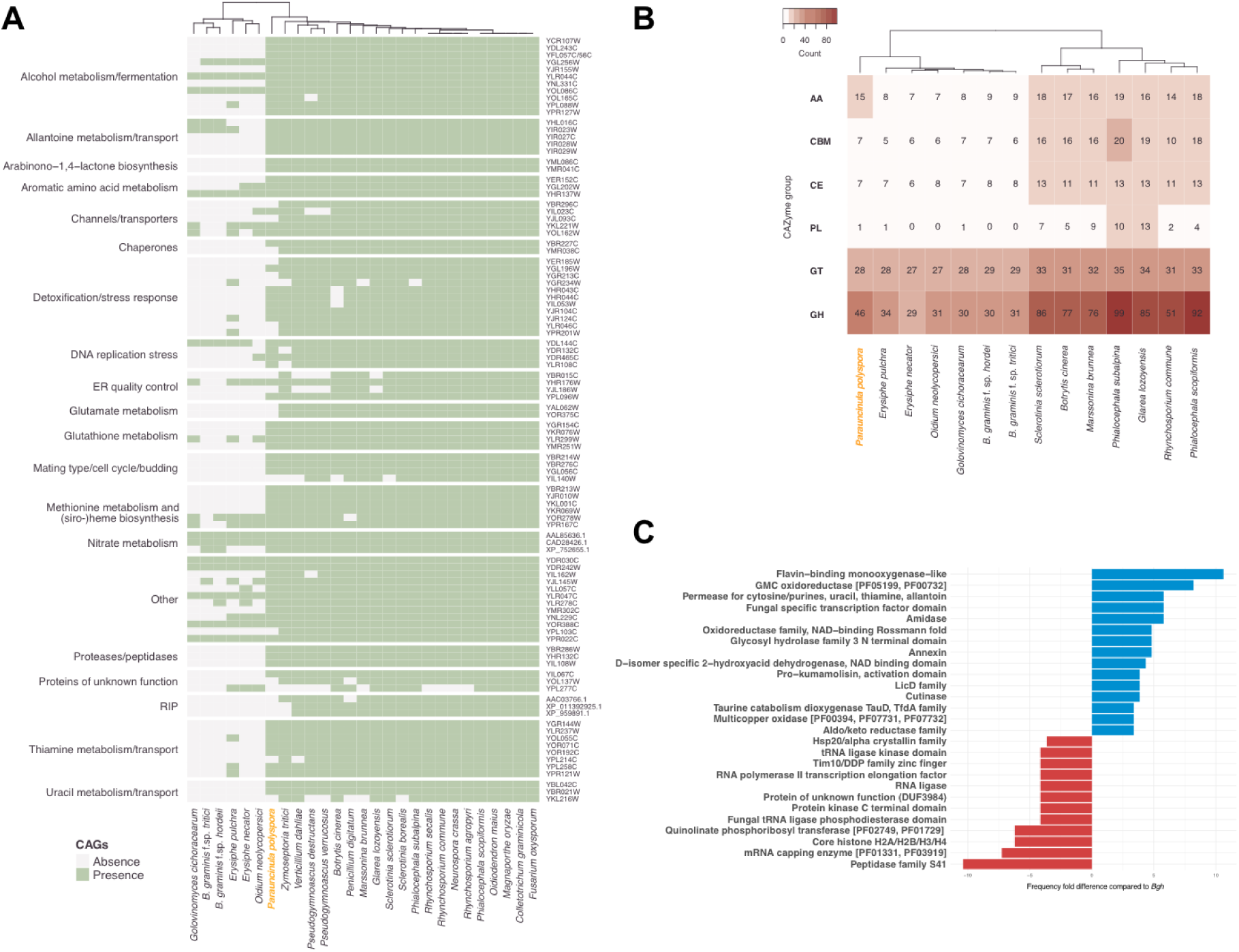
Presence and absence of conserved ascomycete genes (CAGs) in the genomes of powdery mildews. A) Presence and absence of CAGs as cataloged before (Spanu et al. 2010). Hierarchical clustering of the respective species is indicated on the top, gene identifiers are given on the right. B) CAZyme content of different Leotiomycete species in comparison to *P. polyspora*. Abbreviations for CAZyme groups given on the right are as follows: AA, auxiliary activities; CBM, carbohydrate-binding modules; CE, carbohydrate esterases; PL, polysaccharide lyases; GT, glycosyl transferases; GH, glycoside hydrolases. In both A and B the cladograms are based on hierarchical clustering. C) Frequency of fold differences in the occurrence of identified PFAM domains in P. polyspora in comparison to Bgh. Cladograms in A and B were generated by hierarchical clustering.

### *P. polyspora* has a compact predicted secretome with a low number of RNase-like secreted proteins

We identified 261 SP candidates out of which 193 had and 68 lacked a PFAM functional annotation. This is a surprisingly low number when compared to *Bgh* (805 SP candidates, 166 with and 639 without PFAM functional annotation), although other PM species that infect dicotyledonous plant species also have more compact secretomes (450-500 predicted SPs; Wu et al. 2018). However, we cannot exclude the possibility that this comparatively low number reflects, in part, the limitations of our annotation pipeline to predict gene models for SPs, which often lack sequence-relatedness to known proteins, in the absence of transcriptomic data. Nevertheless, out of these 261 predicted SPs only 70 lacked homology to annotated proteins in other PM genomes and merely three (two lacking a PFAM functional annotation) appear to be unique to *P. polyspora*. Regardless if a known domain could be identified, several *P. polyspora* SPs have a homolog in *S. sclerotiorum* and *B. cinerea*, but not in other PMs (Figure 4A), suggesting that these secretome members are dispensable and/or were lost in the course of the adaptation of PM fungi to new hosts. Notably, a sialidase domain-harboring SP (PARAU_11535; homologous to BLGH_03611) seems to be exclusively shared between PMs and appears to be absent in other Leotiomycetes, suggesting this could be an overall conserved virulence-related protein in PMs. Genes encoding ribonuclease (RNase)-like candidate effector proteins are present in high number in the *Bgh* genome (Frantzeskakis et al. 2018; Pedersen et al. 2012). We thus investigated if the predicted secretome of *P. polyspora* likewise contains RNase-like domain-carrying proteins (InterPro accessions: SSF53933, PF06479, PF00445) and whether there is a potential phylogenetic relationship of such proteins with respective homologs in other PMs. We identified only two proteins carrying a recognizable RNase-related domain, which is a surprisingly low number compared to *Bgh* (86 members, Frantzeskakis et al. 2018). We additionally searched for gene models that were excluded from the final annotation (ab initio unsupported calls), and we were able to identify 13 additional genes coding for RNase-like SPs (domain accession SSF53933), which again is still a substantially lower number than in *Bgh*. In the other non-*Blumeria* PM species a similarly low number of secreted RNase-like proteins can be identified (3-19 members, Suppl. Table 8), which suggests that there might be a *Blumeria*-specific expansion of this gene/protein family. Interestingly, after performing ortholog clustering on the mature (signal peptide removed) peptide sequences of the SPs, additional candidate SPs were found to exhibit sequence similarity to these proteins, despite not having a recognizable RNase-like domain (Suppl. Figure 8). In nearly all PM species examined here, family-specific expansions of genes encoding RNase-like SPs can be observed, and in addition, these RNase-like proteins are spread throughout different orthogroups (Suppl. Figure 8), suggesting a polyphyletic origin. We generated a maximum likelihood phylogenetic tree using all PM SP candidates with no PFAM domain (Figure 4B), which further supports the notion that these RNase-like SPs are very diverse and potentially not of monophyletic origin. However, it has been recently suggested that despite the fact that they exhibit a severely eroded primary amino acid sequence similarity, the respective proteins may share a common ancestor, as evidenced by a conserved intron position in the respective genes (Spanu 2017).

**Fig. 4.**
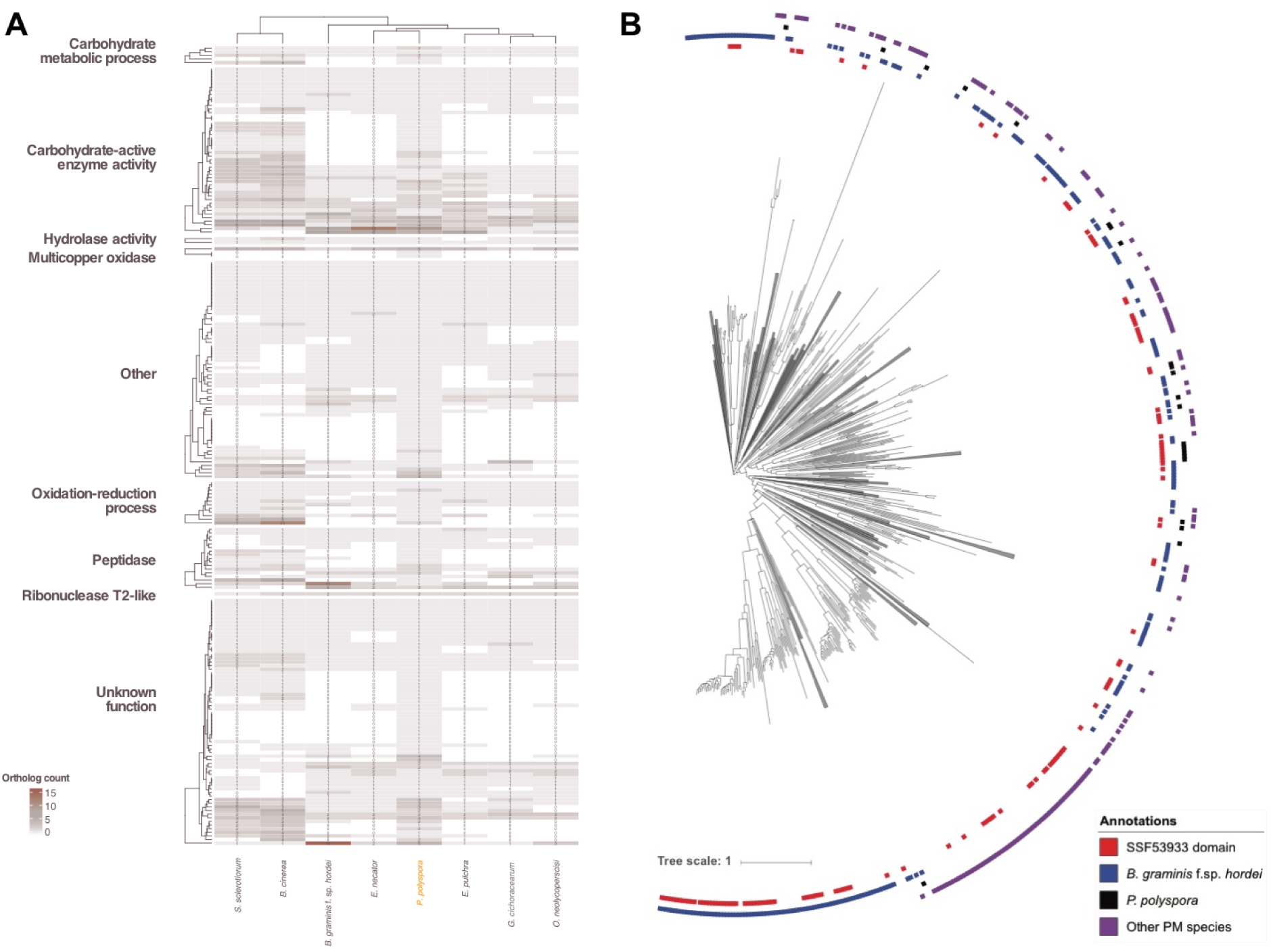
A) Heatmap depicting the differences in the SP content of the publicly accessible powdery genomes and the two related Leotiomycete species *S. sclerotiorium* and *B. cinerea* in comparison to *P. polyspora* according to the color code shown on the left. Description of the functional category of the orthogroups is based on PFAM. The cladograms are the result of hierarchical clustering. B) Maximum likelihood phylogenetic tree of 1227 putative SPs with no PFAM annotation from the proteomes of *Bgh*, *P. polyspora*, *E. necator, E. pulchra, O. neolycopersici* and *G. cichoracearum*. Branches that do not contain and RNase-like domain containing proteins (SUPERFAMILY SSF53933) were collapsed. The category “other PM species” indicated by gray boxes include *E. necator, E. pulchra, O. neolycopersici* and *G. cichoracearum*.

## Discussion

### Co-evolutionary pace and life cycle attributes might drive genome plasticity in PMs

In the work presented here, we attempt to provide a deeper insight in the evolution of the genomic architecture of PM fungi by reconstructing the genome of the early-diverged species *P. polyspora*. Little is currently known about the biology of this fungal pathogen, which seems to propagate exclusively via ascospores produced during sexual reproduction, lacking a recognized asexual morph, i.e. conidiophores and conidia (Meeboon et al. 2017; Takamatsu, Limkaisang et al. 2005), and thus, the asexual part of the typical PM life cycle. Accordingly, its precise host range, mode of infection, the duration of its life cycle and whether it represents indeed a homo- or heterothallic species remain to be explored. With 29 Mb the genome of this species is approximately 4-fold smaller than the average genome of other sequenced PM fungi (Frantzeskakis et al. 2018; Jones, L. et al. 2014; Müller et al. 2018; Wu et al. 2018) and even smaller than that of the average filamentous ascomycete genome (37 Mb; Mohanta and Bae 2015). Accordingly, it lacks the distinct abundance of repetitive elements that otherwise characterizes genomes of PM species (Suppl. Table 5, Figure 2D), but shares with them a comparatively low gene number (6000) and a non-compartmentalized organization (Figure 1C, Suppl. Figure 3; Frantzeskakis et al. 2018). Notably, the genome is indicative of homothallism (self-fertilization) since we located characteristic genes of both mating types (MAT1-1 and MAT1-2) on a single contig (Suppl. Figure 4). This finding is surprising since the majority of PMs is considered to be heterothallic (Tollenaere and Laine 2013), and therefore we expected that our sampling material containing sexual reproduction structures (chasmothecia) should recover discrete scaffolds representing both mating type loci. Nevertheless, homothallism in PM fungi has been reported several times (Tollenaere and Laine 2013), although most of these reports lack molecular evidence. In contrast to the described homothallic *Plantago lanceolata* pathogen *Podosphaera plantaginis*, where both MAT idiomorphs likely exist as functional genes within the same genome (Tollenaere and Laine 2013), in *P. polyspora* the recovered MAT locus contains a seemingly intact MAT1-2-1 and an apparently pseudogenized MAT1-1-3 (Suppl. Figure 4). Moreover, a homolog of MAT1-1-1, supposed to be an indispensable feature of the MAT1-1 mating type (Brewer et al. 2011), seems to be completely absent. Thus, in *P. polyspora* a joint MAT1-1/MAT1-2 locus exists, but it appears as if one of the two idiomoprhs (MAT1-1) became dispensable for sexual reproduction of this fungus. This scenario, where only one MAT idiomorph is sufficient for self-fertility, has been observed previously in ascomycetes (“same-sex mating”; Alby and Bennett 2011), yet it is considered a rare occurrence (Billiard et al. 2011). As in other homothallic ascomycetes and their closely related heterothallic counterparts (Yun et al. 1999), synteny between the *P. polyspora* MAT1-1/MAT1-2 locus and the MAT loci of the closely related heterothallic *Bgh* is poorly conserved. This reported lack of synteny in homothallic loci compared to closely related heterothallic species might also explain why the arrangement of the *P. polyspora* locus is different from the suggested locus of *P. plantaginis* (Tollenaere and Laine 2013). Unfortunately, the fragmentation of the *P. polyspora* assembly and the absence of genomic information on other homothallic PM species precludes further conclusions on the evolution of the mating system in PM fungi at this time. Interestingly, the *P. polyspora* genome retained many genes that were lost in more recently evolved PMs, but similarly to the latter lacks genes associated with the RIP genome defense mechanism (Figure 3A, Figure 2A-C). This finding suggests that the RIP pathway was abandoned in an early progenitor species of the PM lineage at least 80-90 million years ago prior to the separation of the *Parauncinula* genus from the other PMs (Takamatsu, Ninomi et al. 2005). The absence of RIP implies that the propagation of TEs in the genomes of the *Erysiphales* might be restrained by other mechanisms and/or suppressed by certain attributes of the *Parauncinula* life cycle. A plausible hypothesis is that the sexual propagation in *Parauncinula* might contribute to the maintenance of a lean genome, in comparison to the mainly asexually propagating species of the genera *Golovinomyces, Blumeria, Podosphaera* and *Erysiphe* (Glawe 2008). This idea would support the overall assumption that sexual recombination can limit the uncontrolled proliferation of TEs while at the same time help spreading beneficial mutations (Schurko et al. 2009). Nevertheless, examples of fungal species lacking sexual propagation, RIP and an abundance of TEs exist (Man et al. 2016), implying that broad generalizations cannot be easily made. On the other hand, it might be argued that the selection pressure in the *P. polyspora - Q. serrata* interaction is comparatively low. In other cases where PMs infect annual, agriculturally relevant hosts, breeding and growing cultivars with novel resistance specificities as well as crop protection measures place massive selection pressure on the pathogen populations. A particularly plastic and rapidly evolving genome (opposed to the case of *P. polyspora*) should be an advantage during adaptation and survival of crop pathogens. *P. polyspora* likely causes monocyclic infections with a much lower propagation rate, since the lack of conidiophores does not allow for the rapid and profuse aerial dispersal of conidia and multiple infection cycles per year, as for example in *B. graminis*. In addition, the host *Q. serrata* has a long, perennial life cycle, leading to a limited ability to evade the pathogen with a rapid turnover of resistance genes. Also, its genetically diverse local populations (Ohsawa et al. 2008) may offer a less selective environment than the hosts in uniform agricultural settings (Burdon et al. 2013; Zhan et al. 2014). In this scenario, both partners (*P. polyspora* and *Q. serrata*) are locked in an arms race, albeit at a much slower pace compared to the interactions between many annual plant species and the respective PMs. This could be echoed by the *P. polyspora* genome where the relative scarcity of TEs does not offer a template for rapid evolution of virulence genes by duplications (Suppl. Table 5), small scale rearrangements or deletions as it has been proposed for *Bgh* and *Bgt* (Frantzeskakis et al. 2018; Müller et al. 2018, 2018; Praz et al. 2017), since this presumably is not a necessity thus far.

### Functional reduction and family expansion is an ongoing process in PMs

The *P. polyspora* genome indicates that more CAGs are present in an ancestral than in derived PM species and also that the respective gene losses were unequal (Figure 3A). Recent studies indicate that this also extends to CAGs not included here (Wu et al. 2018), as for example the absence of a part of the RNAi machinery in the grapevine PM pathogen *E. necator* (Kusch et al. 2018). Since the remaining CAGs are dispersed in the genome, it is unlikely that their absence is the result of some large-scale genome reduction, e.g. the loss of a single chromosome. Instead, it seems to be in part stochastic, possibly due to local illegitimate recombination activity caused by TE insertions adjacent to the genes (Praz et al. 2017). The eventual fixation of these losses as it is observed in the different PM lineages could be a driver of the subsequent strict association of these species with a low number of specific hosts, and their spatial and reproductive isolation from other PMs. It may also favor the succeeding loss of additional functional categories (Figure 3B, C, Figure 4A) that are related to virulence (i.e. peptidases, CAZymes, redox enzymes) and are no longer needed to that extent in a given highly specialized host environment. This might also apply to asexual reproduction (conidiospore formation), which appears to be absent in species of the genera *Parauncinula, Brasiliomyces* and *Typhulochaeta* within the *Erysiphales* (Glawe 2008). Taken together, the pattern of present/absent genes in the PM genomes thus appears to result from a mixture of divergent, convergent and individual gene losses. We hypothesize that in PMs, extinction by genome erosion is avoided by compensatory lineage-specific expansion of gene families that support virulence (e.g. encoding effector proteins) or other physiological processes. Two examples that were identified here are secreted RNase-like proteins (Figure 4B, Suppl. Figure 7) and Sgk2-type serine/threonine protein kinases (Suppl. Figure 6), which appear to have specifically multiplied their numbers in Bgh. Some RNase-like SPs have been previously shown to be involved in virulence and/or to be recognized by the plant host (Lu et al. 2016; Pennington et al. 2016; Praz et al. 2017; Spanu 2017), while the role of Sgk2-type serine/threonine protein kinases in PM biology remains elusive (Kusch et al. 2014). Yet in other fungal pathogens, it is speculated that kineome expansion is related to environmental and stress responses (DeIulio et al. 2018).

## Conclusions

This study revealed insights in the genome of an early-diverged PM species, thereby providing clues on the evolutionary history of this group of highly host-specialized phytopathogens. Based on the data available so far, we hypothesize the following scenario, which is also summarized in Figure 5. An ancestor of the PM lineage experienced the loss of the RIP machinery and a limited loss of CAGs. After diverging from the other PM lineages, *P. polyspora* experienced a loss of the asexual life cycle, the establishment of homothallism and the expansion of particular protein families (e.g. monooxygenases and oxidoreductases).

**Fig. 5.**
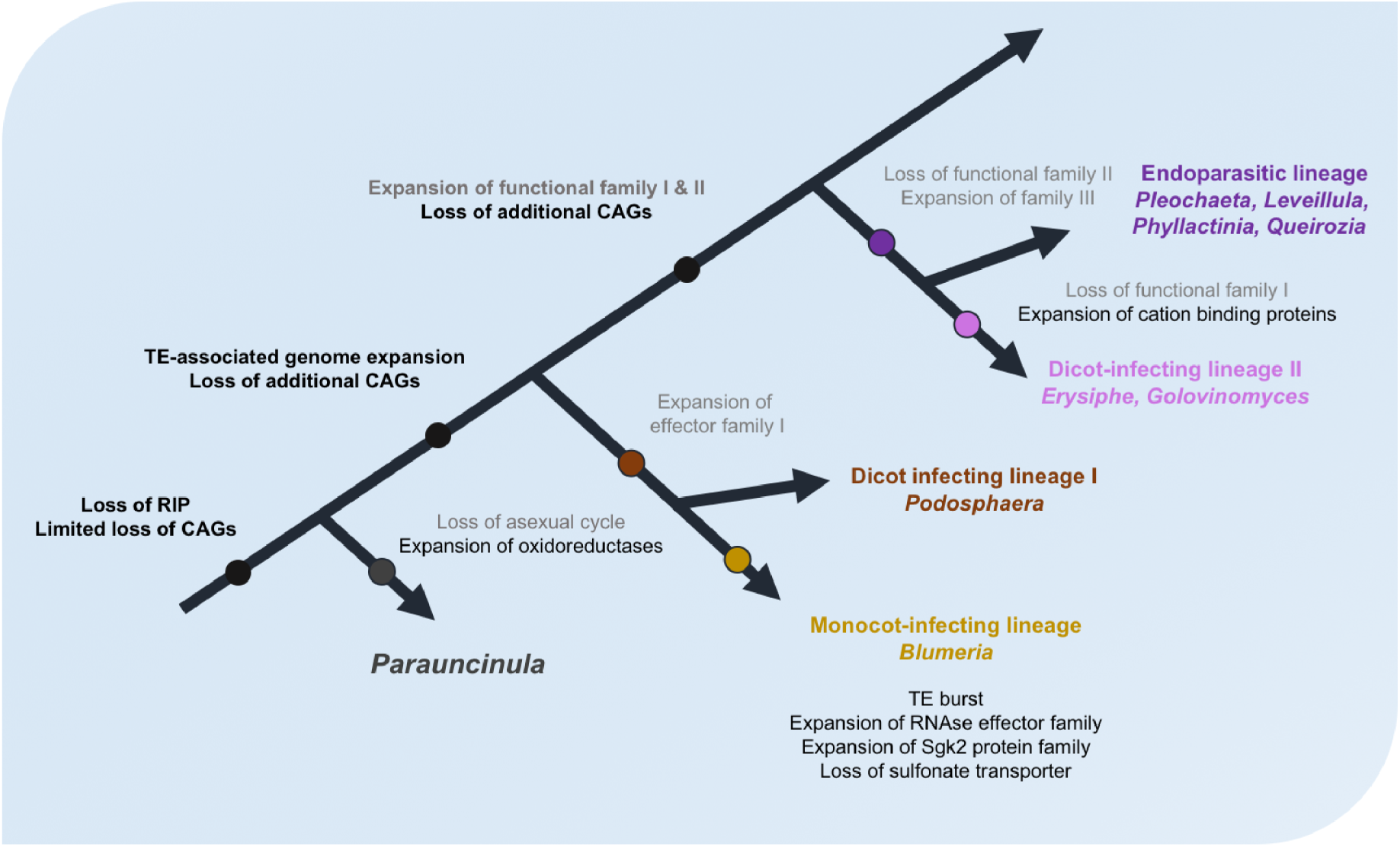
A hypothetical model for the evolution of the PM fungi. A simplified phylogenetic tree illustrating the evolution of PM fungi is shown. Genomic features for which some evidence is provided by the current work and previous studies are shown in black, while hypothetical losses and expansions are depicted in gray. Hypothetical time-points for the events are indicated by circles on the tree, while major/driving events for adaptation to new hosts are highlighted in bold.

The other PM genomes, derived after this early split, underwent a TE-associated genome expansion and the loss of additional CAGs. Separate lineages strictly associated with certain monocot or dicot hosts were established and different modes of infection (epiphytic vs. endophytic) evolved based on a reoccurring pattern of loss and/or expansion of virulence-related and other gene families. This pattern is exemplified by the specific expansion of RNase-like effector proteins and Sgk2 kinases in the monocot-infecting lineage, but also by the unequal loss of CAGs and the reported expansion of other functional families in dicot-infecting PMs (e.g. cation binding proteins; Wu et al. 2018). The genomes of *P. polyspora* and *B. graminis* provide a paradigm of how genome expansion driven by repetitive elements can influence the virulence and the overall biology of fungal plant pathogens. Our study demonstrates that although an excess of TEs might generate genomic architectures that promote rapid adaptability (Dong et al. 2015), this might come with expensive trade-offs such as the loss of critical genes and subsequent (reproductive) isolation in a specialized host niche. This is likely to be setting these plastic genomes in a position where the loss of conserved genes has to be constantly balanced by the expansion of virulence-related gene families. Looking forward, considering the vast range in size and TE content between the different PM genomes, dense sequencing of additional species should provide evidence for progression and different steps in the genome evolution of these rapidly adapting fungal plant pathogens. It would be likewise highly interesting to study the closest saprophytic relatives of the PM lineage, e.g. members of the *Myxotrichaceae* (Takamatsu 2013) at the genomic level. Such comparative analyses promise to unravel genomic adaptations that enabled the transition from a saprophytic to a pathogenic way of life, and ultimately the shift to an obligate biotrophic lifestyle. The comparative analysis of the *P. polyspora* genome with that of other PM species may further reveal gene sets that are necessary for an asexual life cycle, i.e. the formation of conidiophores and the production of conidia.

## Materials and Methods

### Sampling and genomic DNA extraction

Infected leaves of a single *P. polyspora* infected *Q. serrata* tree were collected in 2017 in Torimiyama Park, Haibara, Uda-shi, Nara Prefecture, Japan (N34.543809, E135.944306). The leaf samples were dipped in 5% w/v cellulose acetate-acetone solution, and then placed to dry. Using forceps, the cellulose was peeled off and the sample ground in liquid nitrogen with mortar and pestle. The resulting cellulose fragments with fungal structures attached were transferred to 2 ml tubes and genomic DNA extraction performed as described in (Feehan et al. 2017). Afterwards, small DNA fragments (<100 bp) were removed using AMPure XP beads (Beckman Coulter, Germany) and the quantity and quality of the DNA was assessed using a NanoDrop spectrophotometer (Thermo Scientific, Germany) and a Qubit fluorometer (Invitrogen, Germany).

### Genome sequencing, assembly and functional annotation

Illumina library preparation (TruSeq DNA Nano, llumina) and genomic sequencing was performed by CeGaT GmbH in Tübingen, Germany. The library was sequenced on the NovaSeq 6000 platform and resulted in 163,1 million paired raw reads (2×150 bp, total of 24,6 Gbp data). The reads were assessed for their content of Leotiomycete sequences using MG-RAST (MG-RAST id: f5bdd547896d676d343830363131372e33, Meyer et al. 2017). The pipeline followed to assemble the genome is briefly presented in Suppl. Figure 1A. In more detail, the adapters were trimmed with Skewer (Jiang et al. 2014), and then passed to BFC (−b 32 −k 25 −t 10, Li 2015) for error correction and removal of singleton kmers. The corrected reads were then assembled with SPAdes v3.11.1 (–only-assembler -k 31,51,71,91,111 –meta, Bankevich et al. 2012). In order to remove bacterial and eukaryotic contaminating sequences from the resulting scaffolds, the sequences were initially searched by BLAST against a set of 3837 plant associated bacterial genomes (http://labs.bio.unc.edu/Dangl/Resources/gfobap_website/index.html; Levy et al. 2018). The resulting scaffolds were filtered based on their depth (cutoff >20x, Suppl. Figure 1C) and the homology of their annotations to the Leotiomycetes. For the exclusion based on the Leotiomycete homology, after the annotation (see below) the predicted genes were used for homology search against the NCBI nr protein database (last accessed 11/2017) using BLAST+ v2.3.0 (Camacho et al. 2009). Scaffolds where the two most frequent hits belonged to the Leotiomycetes were deemed as high confidence scaffolds, while the rest were rejected. These high confidence scaffolds were assessed for coverage of the gene space using BUSCO v1.22 (Simão et al. 2015), and an additional size estimation based on kmer abundance was provided using Jellyfish v2.2.10 (Marçais and Kingsford 2011). For the annotation of the scaffolds we followed the same pipeline as described before (Frantzeskakis et al. 2018) using MAKER (Holt and Yandell 2011). The datasets provided as evidence are listed in Suppl. Table 9. Afterwards functional annotation was performed using InterProScan v5.19-58.0 (Jones, P. et al. 2014), and HMMER v3.1 (Finn et al. 2011) with dbCAN v6 (Le Huang et al. 2018) for the identification of CAZymes specifically. Putatively secreted proteins with no transmembrane domains were identified using SignalP v4.1 (Petersen et al. 2011) and TMHMM v2.0c (Emanuelsson et al. 2007). Mating type genes were identified using bi-directional BLAST searches (Camacho et al. 2009) using BLASTP and TBLASTN with an evalue cut-off of 10e-5.

### Analysis of repetitive sequences

Repetitive se-quences were identified using RepeatMasker v4.0.6 (www.repeatmasker.org) using Repbase as a database (last accessed 2016/06/09). Subsequently, a repeat landscape was generated for P. polyspora as described before (Frantzeskakis et al. 2018). GC composition of the selected Leotiomycete genomes and dinucleotide frequencies were calculated using OcculterCut v1.1 (Testa et al. 2016) and RIPCAL v2 (Hane and Oliver 2008), respectively.

### Orthogroup inference, phylogeny and nucleotide polymorphisms

Identification of ortholog groups and generation of gene family trees was performed using OrthoFinder v1.1.2 (Emms and Kelly 2015). The maximum likelihood phylogenetic trees based on putatively secreted proteins with no PFAM annotation or on single copy orthologs, were generated using FastTree v2.1.10 (Price et al. 2010) after alignment of the protein sequences with MAFFT v7.310 (Katoh and Standley 2013). Figures of the trees were generated using iTOL (Letunic and Bork 2016). CAG search using the proteomes listed in Table S9 was performed using BLASTP (Camacho et al. 2009) with an e-value threshold of 10e-5. In order to discover the number of single nucleotide polymorphisms in the P. polyspora genome assembly we initially mapped the reads using BWA-MEM v0.7.15-r1140 (Li 2013). The resulting sam file was processed (conversion to bam, sorting) with Picard tools v2.8.2 (http://broadinstitute.github.io/picard), and then polymorphisms were identified using samtools mpileup and bcftools (v0.1.19, Li et al. 2009) and filtered with SnpSift v4.3i (QUAL >= 20 && DP > 3 && MQ > 50, Cingolani et al. 2012).

## Code and data availability

Corresponding R scripts and associated files (phylogenetic trees, tables, etc.) used for generating figures used in the paper are deposited in https://github.com/lambros-f/paraun_2018. The dataset, including raw reads and the assembled genome used here are deposited on ENA under the accession number PRJEB29715.

## Supporting information

## Acknowledgements

Diana Seress’ contribution to sample preparation is acknowledged. This work was supported by a grant of the Deutsche Forschungsgemeinschaft (DFG)-funded Priority Programme SPP1819 (Rapid evolutionary adaptation - Potential and constraints) to R.P. (PA 861/14–1). The analysis was performed with computing resources granted by RWTH Aachen University under project rwth0146.

## Author contributions

RP and LF conceived the study and drafted the manuscript. ST did the field sampling, MZN and LK contributed to sample preparations. LF and MZN performed the experiments. LF analyzed the data. LF, MB and SK analyzed the mating loci. RP, ST and LK provided conceptual advice. All authors have edited, read and approved the final manuscript version.

## Supplementary Figures

**Suppl. Figure 1:**
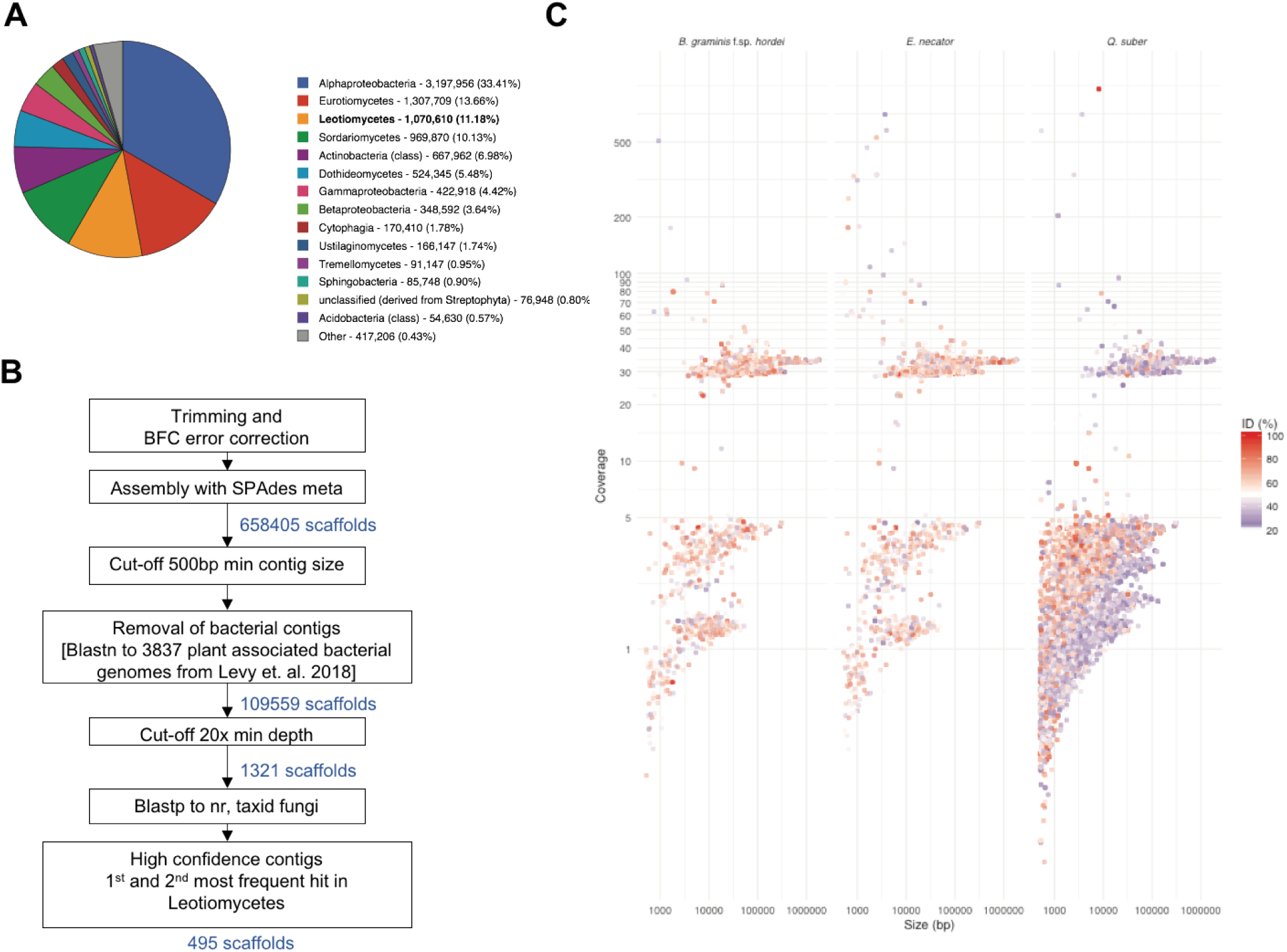
Bioinformatic pipeline and quality control of the assembled sequences.A) Taxonomic hits distribution based on the MG-RAST analysis by class. Hits with lower than 0.5% distribution were group as “Other”. B) Assembly pipeline for the *P. polyspora* genome. C) TBLASTN matches of coding sequences contained in the pre-filtered *P. polyspora* scaffolds, to the genomes of two powdery mildew species (*Bgh* and *E. necator*) and an oak species (*Q. suber*) closely related to the host of *P. polyspora*. Coverage is derived by the metaSPAdes result.

**Suppl. Figure 2:**
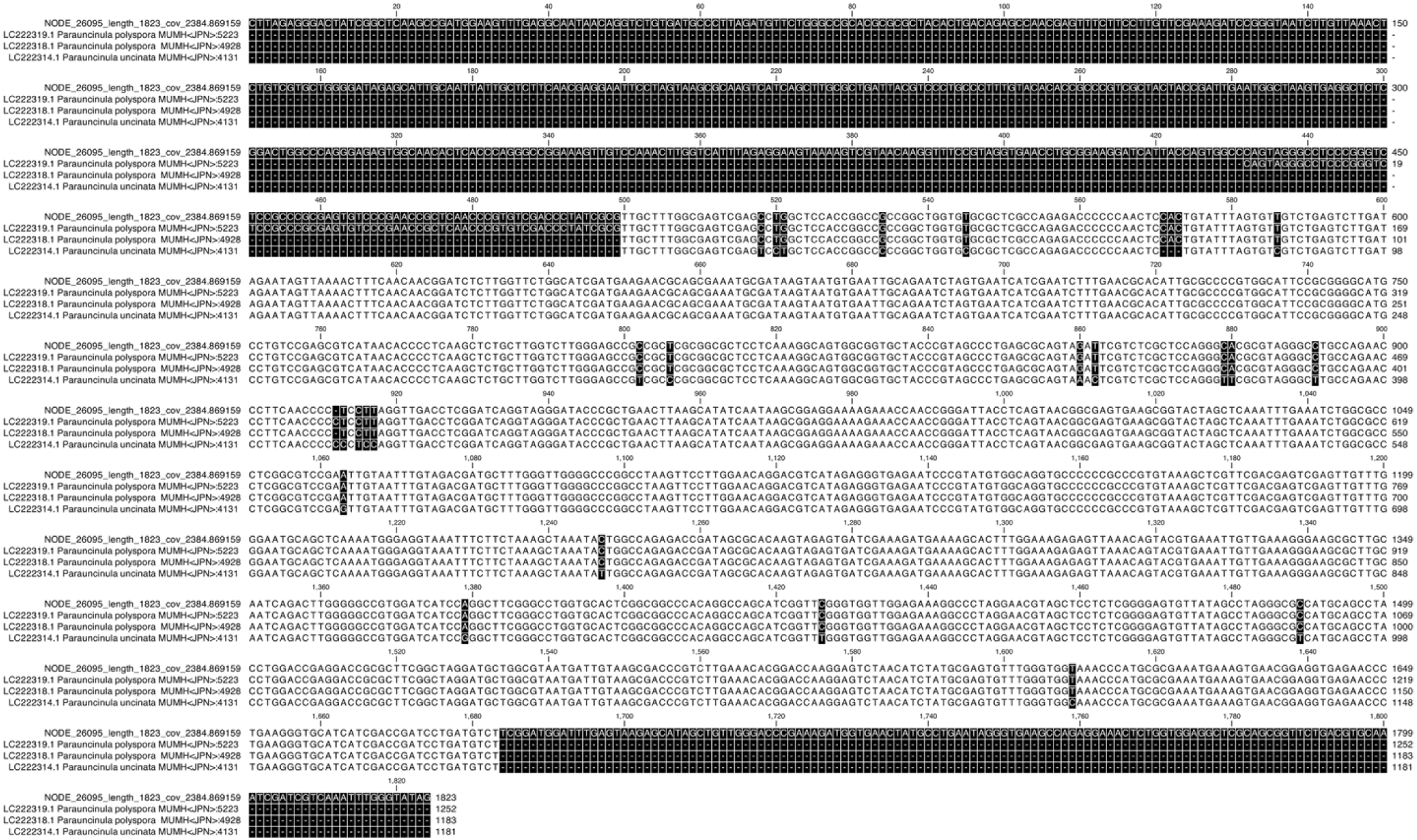
Multiple sequence alignment of an ITS sequence-harboring scaffold with known ITS sequences of the genus *Parauncinula*. The multiple sequence alignment was performed with CLC Sequence Viewer (QIAGEN) and comprises the assembled scaffold “NODE_26095_length_1823_cov_2384.869159” and the reference ITS sequences for *P. polyspora* and the related species *P. uncinata*.

**Suppl. Figure 3:**
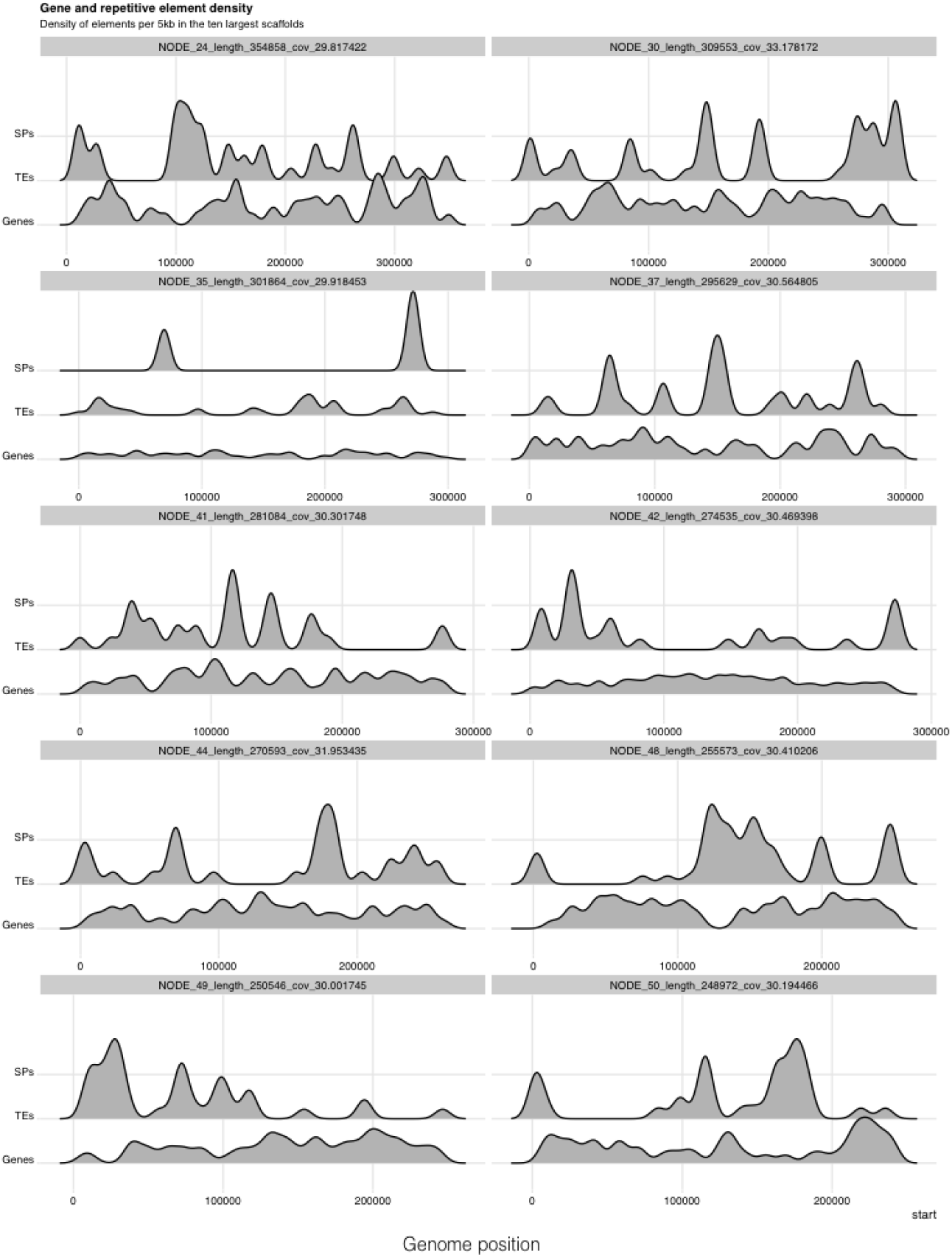
Distribution and density of different genetic elements in the *P. polyspora* genome. Relative density of genes encoding putative SPs, non-SP-coding genes and TEs in the ten largest scaffolds of the *P. polyspora* genome.

**Suppl. Figure 4:**
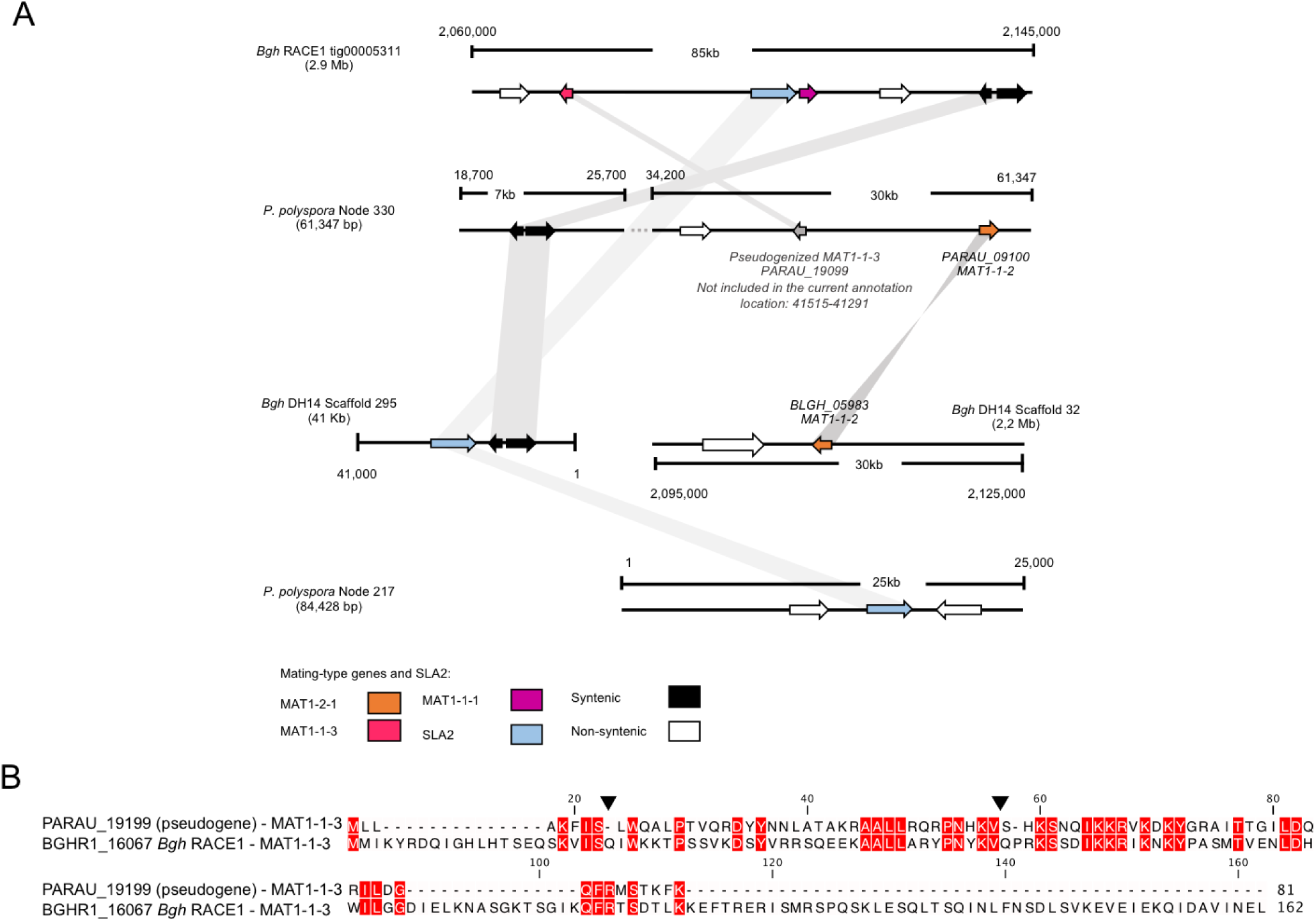
Analysis of the *P. polyspora* mating type locus. A) Schematic representation of the MAT loci in *P. polyspora* and *B. graminis* f.sp. *hordei*. MAT-associated genes are colored while flanking genes which are homologous and have a degree of synteny between the two species are shown in black. White arrows depict genes that are non-syntenic but homologous. B) Alignment of the putative *P. polyspora* MAT1-1-3 pseudogene and its *Bgh* homolog (BGHR1_16067). Black arrowheads point the positions of stop codons in the *P. polyspora* sequence. Sequence conservation (100% identity) is highlighted in red.

**Suppl. Figure 5:**
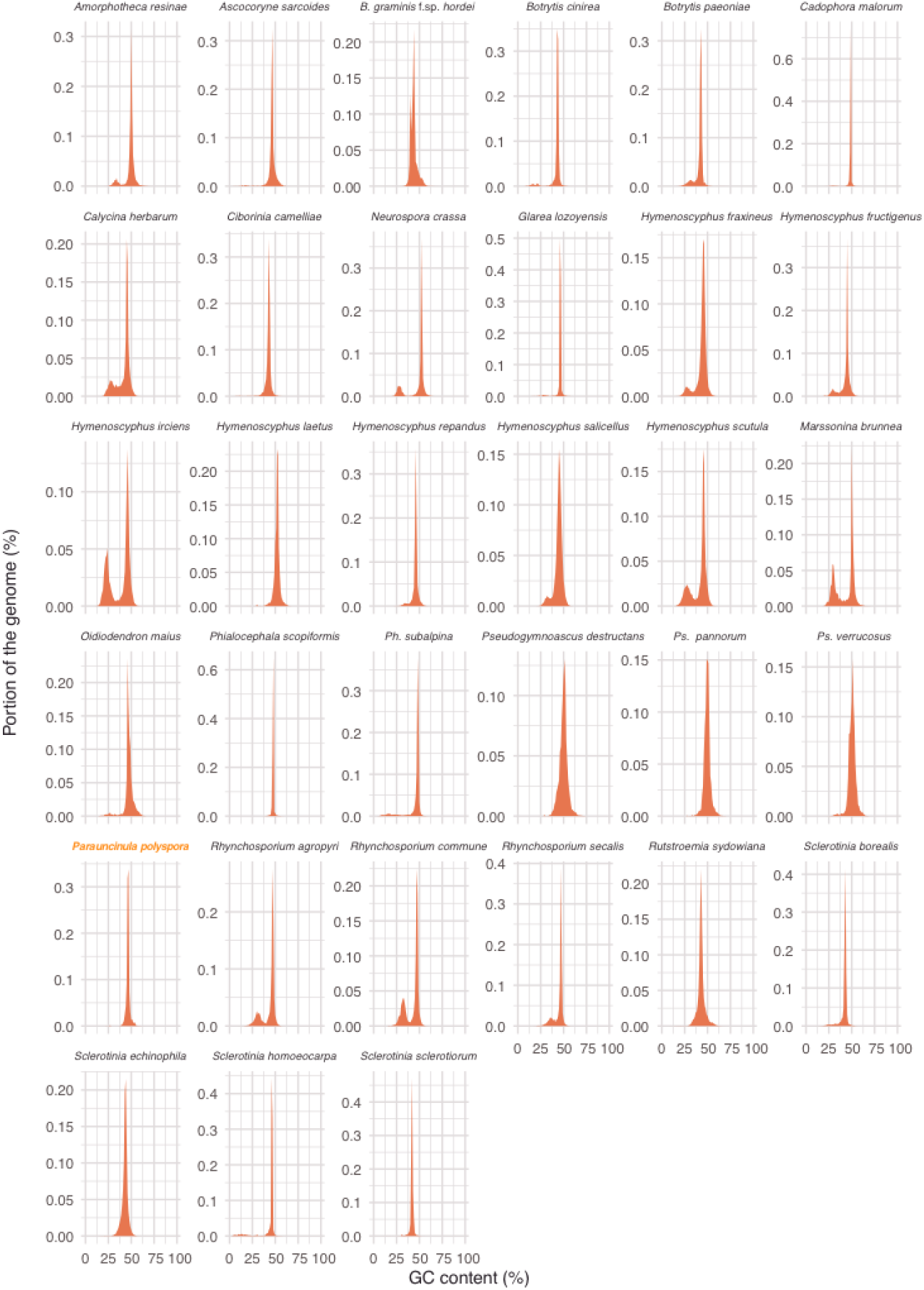
Presence/absence of AT isochores in the Leotiomycetes. Thirty-three Leotiomycete genomes were assayed for the presence of AT isochores using OcculterCut v1.1. The portion of the genome with the respective % GC content are plotted indicating the presence of AT isochores in cases where a bimodal distribution is present.

**Suppl. Figure 6:**
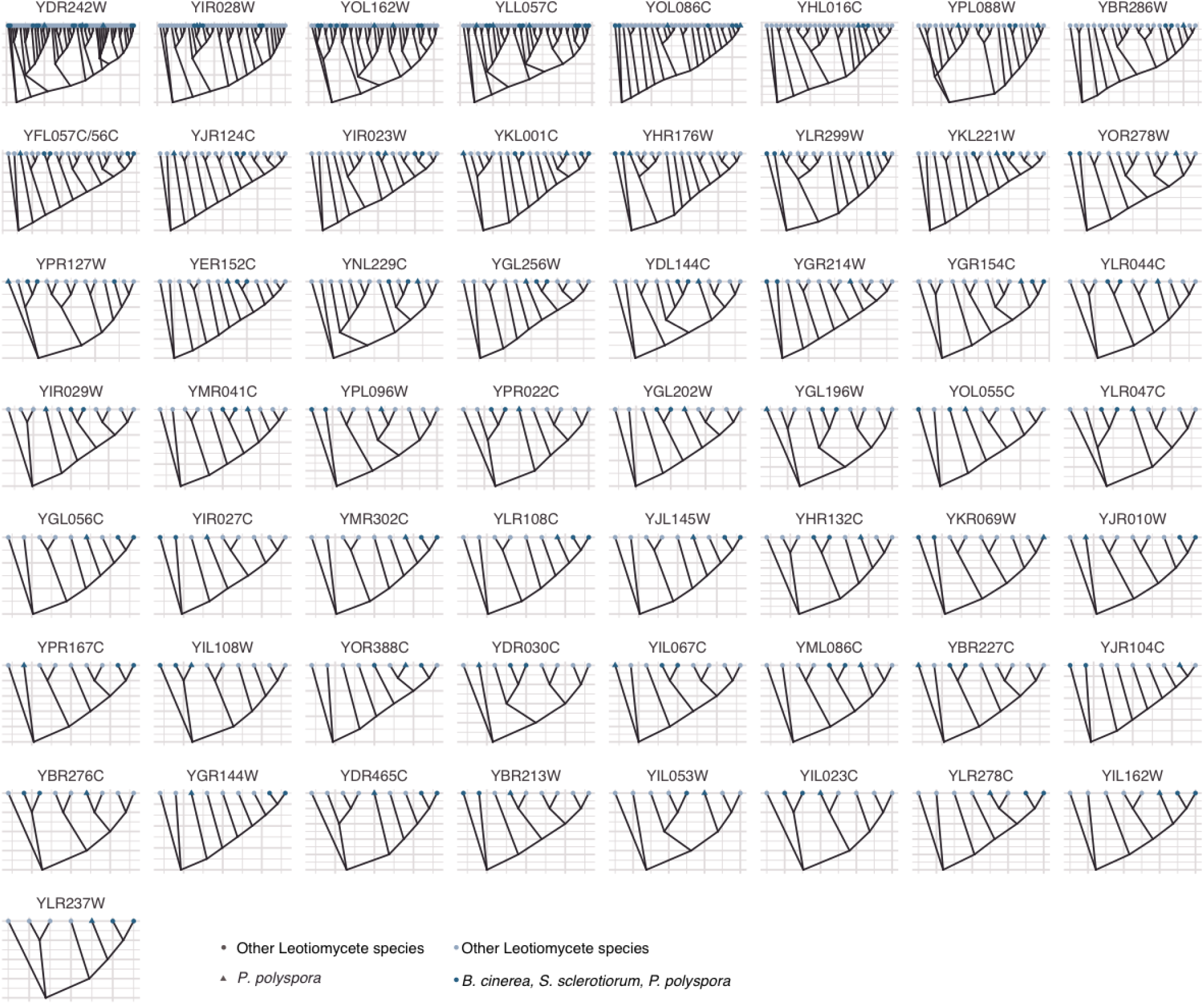
Maximum likelihood phylogenetic trees per CAG orthogroup. For every conserved CAG for which a *P. polyspora* homolog exists, a phylogenetic tree generated by Orthofinder and drawn to assess the position of the homolog in relation to the species phylogeny presented in Figure 1A (for those species having a respective ortholog).

**Suppl. Figure 7:**
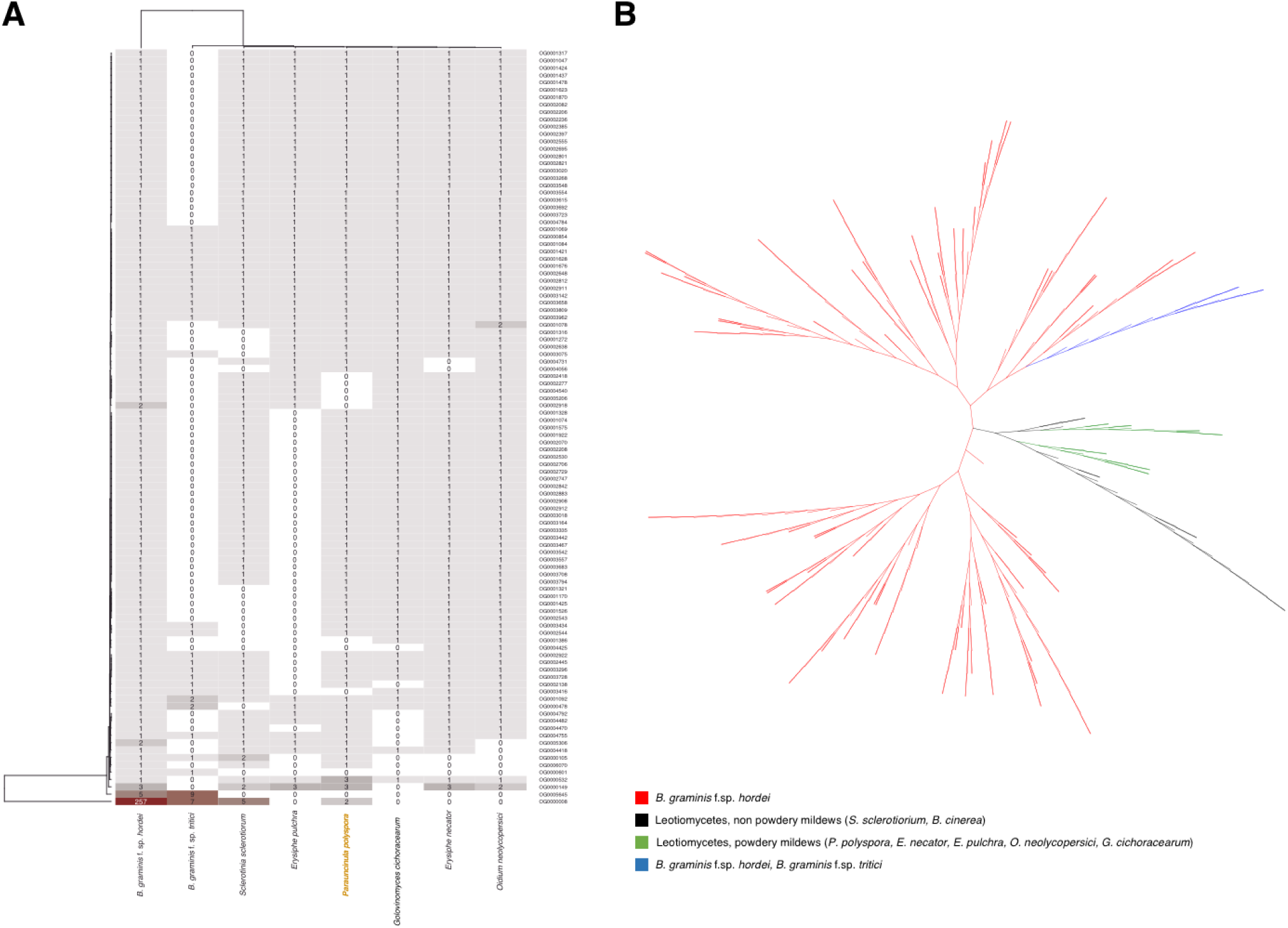
Expansion of Sgk2-type serine-threonine/tyrosine-protein kinases in *Blumeria*. A) Heatmap of orthogroups containing proteins with the Sgk2 domain (SUPERFAMILY SSF56112). B) Maximum likelihood phylogenetic tree with Leotiomycete Sgk2 members. Red highlights branches with *Bgh* members only, blue indicates *Bgh* and *Bgt*, green indicates *P. polyspora* and other powdery mildew species, while black indicates other Leotiomycete species.

**Suppl. Figure 8:**
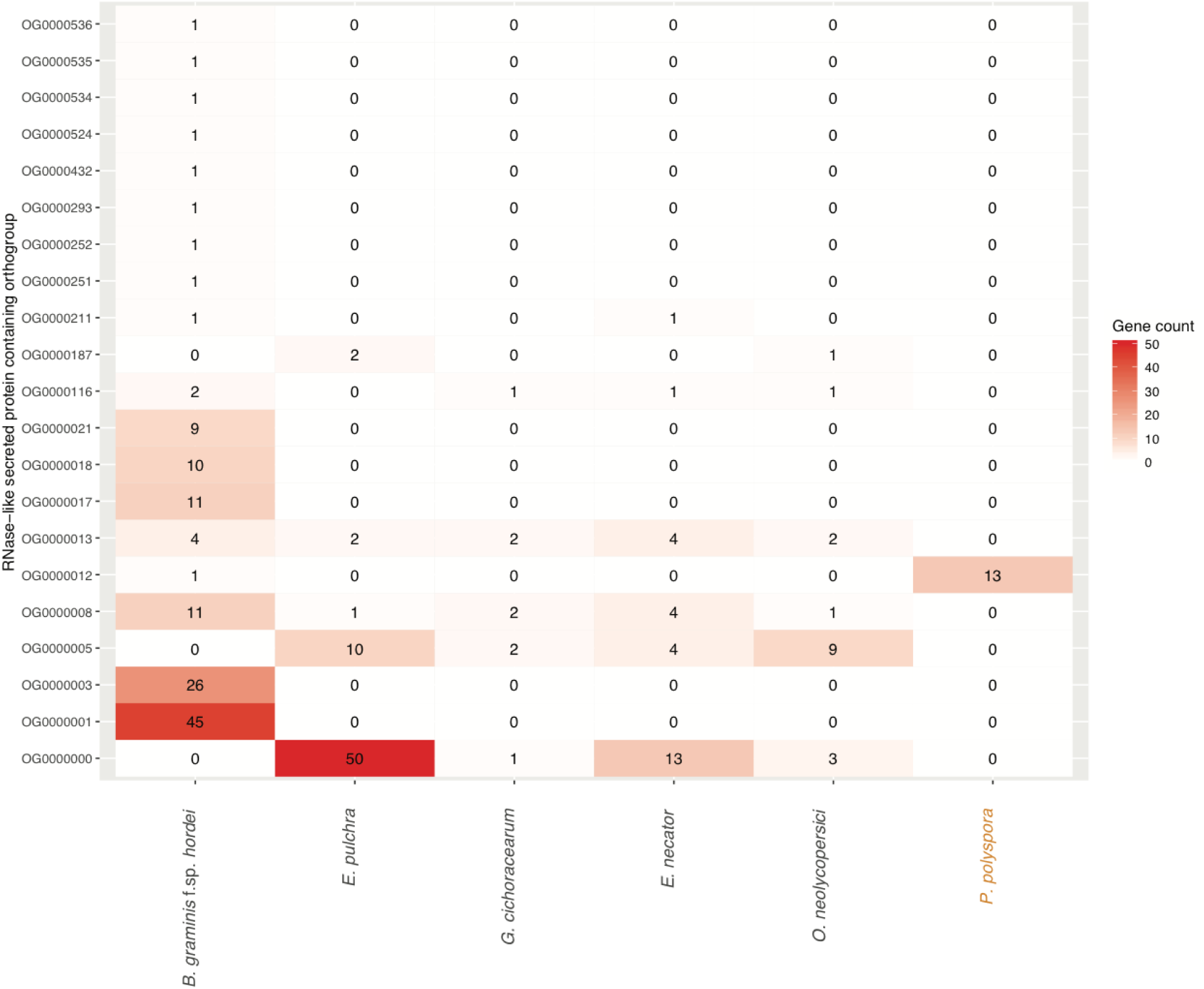
Orthogroup clustering of RNase-like SPs. Predicted powdery mildew SPs were clustered based on their mature (signal peptide removed) peptide sequence using Orthofinder. The heatmap depicts the abundance of proteins for every orthogroup containing at least one SP with an RNase/RNase-like domain (SSF53933). Color code is shown on the right.

